# Characterization of an Open-Channel Structure and Lateral Conduction Pathway in the Cation-Selective Pentameric Ligand-Gated Ion Channel, ELIC

**DOI:** 10.1101/2025.11.06.686614

**Authors:** Mark J. Arcario, Elizabeth J. Wu-Chen, Jérôme Hénin, Grace Brannigan, Wayland W.L. Cheng

**Affiliations:** Department of Anesthesiology, School of Medicine, Washington University in Saint Louis, St. Louis, MO, 63110, USA; Laboratoire de Biochimie Théorique, Université Paris Cité, Paris, 75005, France; Center for Computational and Integrative Biology, Rutgers University, Camden, NJ, 08103, USA; Department of Physics, Rutgers University, Camden, NJ, 08102, USA

## Abstract

Open-channel structures of multiple pentameric ligand-gated ion channels (pLGIC) have been determined, including the prokaryotic model pLGIC, ELIC (*Erwinia* ligand-gated ion channel). For many of these structures, it remains uncertain whether they represent the physiologic open-channel state because the conditions used for structure determination do not match those of functional measurements in cell membranes. Here, molecular dynamics (MD) simulation is used to examine the ion conduction properties of the ELIC open-channel structure, which was determined using a non-desensitizing mutant called ELIC5. Results from simulations show that the pore remains stably open on the microsecond timescale, but computational electrophysiology measurements demonstrate a large outward rectification, and an inward conductance that is significantly lower than experiment. This discrepancy is attributed to a constricted extracellular domain (ECD), which restricts the passage of ions between the ECD vestibule and extracellular solution. Unbiased MD simulation of the ELIC5 structure demonstrates spontaneous widening of an intersubunit space in the ECD to expose a lateral fenestration which becomes the dominant ion conduction pathway. Computational electrophysiology of the ELIC5 MD-refined structures with a widened lateral fenestration shows better agreement with experimental single-channel recordings. Mutations of residues along the lateral ion conduction pathway show reduced single-channel conductance, supporting the importance of the lateral fenestration for ion conduction in a cation-selective pLGIC.

## Introduction

Structural biology approaches, including X-ray crystallography (XRC) and cryo-electron microscopy (cryo-EM), provide complementary insights to biochemical and electrophysiological studies in understanding the complex gating, allostery, and ion permeation of pentameric ligand-gated ion channels (pLGICs). The number of pLGIC structures has increased exponentially with advances in single particle cryo-EM and membrane mimetic platforms (Howard, 2021). Despite advances in methodology, capturing conductive or open-channel structures of pLGICs using XRC and cryo-EM remains challenging. When activated, these channels undergo conformational changes to a metastable, short-lived open-channel state (lasting on the order of milliseconds) before transitioning to the more stable desensitized state. Not only is the open-channel state transient, but the conditions for structural characterization of pLGICs, whether via crystallization or cryogenic temperatures, differ significantly from the conditions used to study their functional properties (Gonzalez-Gutierrez et al., 2017).

Since the description of a purported open-channel state of *Torpedo* nAChR (at 9 Å resolution) (Unwin, 1995), structures of other pLGICs have been reported in apparent open-channel conformations (Table 1; GLIC: Bharambe et al. (2024); Bocquet et al. (2009); Sauguet et al. (2013), ELIC: Petroff II et al. (2022), sTeLIC: Hu et al. (2018), DeCLIC: Hu et al. (2020), *C. elegans* GluCl: Hibbs and Gouaux (2011), human *α*7 nAChR: Noviello et al. (2021), mouse 5HT_3A_: Basak et al. (2018), zebrafish *α*1 GlyR: Du et al. (2015); Kumar et al. (2020), GABA_A_ Mihaylov et al. (2025)). Strategies such as time-resolved freezing (Mihaylov et al., 2025; Unwin, 1995), allosteric modifiers (Hibbs and Gouaux, 2011; Noviello et al., 2021), pore blockers (Kumar et al., 2020), and mutations (Gonzalez-Gutierrez et al., 2012, 2017; Petroff II et al., 2022; Sauguet et al., 2013) have been used to increase the probability of the channel being in the open-channel state. In contrast, some pLGIC structures show widely open-channel pores in the presence of agonist alone (Bocquet et al., 2009; Hu et al., 2020, 2018), where the majority of channels are expected to be in a desensitized state with a narrow, non-conducting pore. Complicating this picture further is the description of a set of ELIC mutants that stabilize the open-channel state in single channel studies, but XRC structures of these mutants in the presence of agonist show no difference from unliganded, resting-state structures (Gonzalez-Gutierrez et al., 2012). Given the discrepancy between the conditions used for cryo-EM of pLGICs and electrophysiology measurements of ion conduction, it remains unclear, in most cases, whether the conformation determined represents the physiologic open-channel state. Indeed, in several cases, pLGIC structures deemed to be “open” have had their TMD pores rapidly collapse and dehydrate in molecular dynamics (MD) simulation (Cerdan et al., 2018; Cheng and Coalson, 2012; Li et al., 2023; Nury et al., 2010).

**Table 1:** Comparing the purported open-channel state structures of pLGICs.

| Channel | PDB<sup>1</sup> | Res.<br>(Å) | Membrane Mimetic | Stabilizing<br>Factors | Min. Pore<br>Diameter<br>(Å) | Computed<br>Conductance (pS)<sup>2</sup> |
| --- | --- | --- | --- | --- | --- | --- |
| DeCLIC | 6V4A | 3.83 | Micelle (DDM) | – | 10.2 | – |
| ELIC5 | 8TWV | 3.40 | spNW15 Nanodisc<br>(POPC, POPE, POPG) | – | 6.6 | <sup>3</sup> |
| ELIC5 | 9NGC | 2.80 | Liposome (POPC, POPE,<br>POPG) | – | 4.4 | – |
| GLIC | 3EAM | 2.90 | Micelle (DDM) | – | 5.0 | – |
| GLIC | 4HFI | 2.40 | Micelle (DDM) | – | 5.0 | – |
| GLIC | 5V6N | 3.35 | Micelle (DDM) | C27S/K33C/<br>I233A/N245C,<br>DTT | 5.0 | 18<sup>4</sup> (9) (Gonzalez-Gutierrez et al., 2017) |
| GluCl | 3RHW | 3.30 | Mixed Micelle (DDM,<br>POPC, DPPC) | anti-GluCl Fab,<br>Ivermectin | 4.6 | – |
| $\alpha 1$ GlyR | 3JAE | 3.90 | Micelle (DDM) | – | 8.8 | 413<sup>3</sup> (80-88) (Cerdan<br>et al., 2018) |
| $\alpha 1$ GlyR | 6UD3 | 3.50 | MSP1E3D1 Nanodisc<br>(Soybean polar extract) | Picrotoxin | 6.5 | 20<sup>5</sup> (80-88) (Kumar<br>et al., 2020) |
| $\alpha 7$ nAChR | 7KOX | 2.70 | Saposin Nanodisc (Soy-<br>bean polar extract,<br>cholesterol) | PNU-120596 | 7.2 | 76<sup>4</sup> (186) (Zhuang<br>et al., 2022) |
| sTeLIC | 6FL9 | 2.30 | Mixed Micelles (DDM,<br>NDG) | – | 11 | – |
| 5-HT<sub>3A</sub> | 6DG8 | 3.89 | Micelle (C<sub>12</sub>E<sub>9</sub>) | – | 6.4 | – |

Commonly, the functional state ascribed to a pLGIC structure is based on the minimal diameter of the transmembrane domain (TMD) pore, specifically if the minimal diameter is large enough to pass the permeant ion. However, pLGIC structures with pore diameters larger than this cutoff may still pose an energetic barrier to ion conduction especially at hydrophobic residues (Basak et al., 2018; Bharambe et al., 2024). In pLGICs known to have one predominant conductance state, the exact pore structure can be affected by the membrane mimetic (Dalal et al., 2024, 2025) or the choice of allosteric modifier (Du et al., 2015; Kumar et al., 2020), suggesting that the pore diameter is not the sole determinant of conductance. Furthermore, the extracellular (ECD) and intracellular (ICD) domains, which are known to affect conductance (Hales et al., 2006; Kelley et al., 2003) and gating (Ivica et al., 2021), are generally neglected when considering the pore profile of pLGIC structures. Perhaps the biggest drawback to this approach is that it ascribes a static structure to a dynamic functional state.

Recently, more dynamic metrics from MD simulations have been used in assigning functional states to pLGICs structures. The use of pore diameter as a metric (Smart et al., 1996) has been adapted to incorporate ellipsoidal shapes (Seiferth and Biggin, 2024), which may more accurately capture the geometry of the pore, as well as measures of pore hydrophobicity and hydration (Klesse et al., 2019). While a hydrated ion conduction pore is likely necessary for ion permeation, hydration is not equivalent to ion permeability.

Ideally, any metric to validate the functional state of a structure would be compared against experimental values. In this regard, Cecchini and colleagues performed computational electrophysiology on the zebrafish *α*1 GlyR and compared the results to experimental single channel ion conductance and polyatomic ion permeability (Cerdan et al., 2018). In this approach, a static electric field is applied across the simulation cell resulting in a fixed transmembrane potential difference (ΔV_m_) (Gumbart et al., 2012). The magnitude of ΔV_m_ can be tuned by altering the strength of the fixed electric field, allowing measurement of single channel currents at multiple potentials. Using this approach, it was shown that the zebrafish *α*1 GlyR purported open-channel structure (PDB 3JAE) was too open (the computational single channel conductance was four times experimental measurements) and that a more constricted pore, similar to other purported open-channel structures (Bocquet et al., 2009; Hibbs and Gouaux, 2011) was more representative of the open-channel state of *α*1 GlyR. The same conclusion was reached in a different study where picrotoxinin was shown to not completely block ion conduction through the open-channel model of *α*1 GlyR (Gonzalez-Gutierrez et al., 2017). The computational electrophysiology approach has since been applied to other pLGICs (Table 1) (Avstrikova et al., 2025; Basak et al., 2018; Cerdan and Cecchini, 2020; Cerdan et al., 2022; Dämgen and Biggin, 2020; Gonzalez-Gutierrez et al., 2017; Zhuang et al., 2022) and similar methods have been introduced which incorporate asymmetric salt concentrations (Khalili-Araghi et al., 2013; Kutzner et al., 2011) or calculate ion permeability coefficients from free energy surfaces (Pohorille and Wilson, 2021). These methods allow direct comparison between the structural data and functional data, increasing confidence in assigned functional states.

An open-channel structure was obtained of ELIC, a prokaryotic pLGIC (*Erwinia* ligand-gated ion channel), using a combination of five mutations (P254G/V261Y/C300S/G319F/I320F, called ELIC5) that eliminate desensitization (Petroff II et al., 2022). ELIC5 is a unique model of the pLGIC open-channel state, with an open probability near 1 at saturating agonist concentration. Here, using MD simulations coupled with functional measurements, we characterize the stability and ion conductance properties of the ELIC5 open-channel structure. While the pore is stably hydrated and apparently open on the microsecond timescale, computational electrophysiology shows an inward conductance that is lower than expected. The simulations reveal a lateral fenestration in the ECD that enlarges and becomes the main pathway for ion conduction after extended equilibrium simulation, similar to the lateral pathway observed in the *α*1 GlyR (Cerdan et al., 2022). Computational electrophysiology of ELIC5 with the enlarged lateral pathway reproduces experimental values of conductance. Mutagenesis of acidic residues in the lateral pathway confirms the role of this pathway in determining the inward conductance of ELIC. Together, these results validate an open-channel conformation of ELIC revealed using unrestrained MD simulations and highlight the importance of the lateral pathway in the conductance of a cation-selective pLGIC.

## Materials and Methods

### Materials

1-palmitoyl-2-oleoyl-*sn*-glycero-3-phosphatidylcholine (POPC), 1-palmitoyl-2-oleoyl-*sn*-glycero-3-phosphatidylethanolamine (POPE), and 1-palmitoyl-2-oleoyl-*sn*-glycero-3-phosphatidylglycerol (POPG) were obtained from Avanti Polar Lipids (Alabaster, AL). BioBeads SM-2 adsorbent beads were purchased from Bio-Rad Laboratories (Hercules, CA). All other chemicals were purchased from Sigma-Aldrich (St. Louis, MO).

### Expression, Mutagenesis, and Purification of ELIC

pET26-MBP-ELIC (gift from Raimund Dutzler, AddGene plasmid #39239) was used for expression of wild-type ELIC and as template DNA for all mutations. All agar plates and liquid growth media contained kanamycin at 50 *µ*g/mL. Point mutations were introduced into the template plasmid DNA with the QuikChange II site-directed mutagenesis kit (Agilent Technologies, Santa Clara, CA) per manufacturer’s protocol with double mutants produced by sequential point mutations. Mutagenesis was confirmed by Sanger sequencing (Genewiz, South Plainfield, NJ). Each plasmid was transformed into OverExpress C43(DE3) competent cells (Biosearch Technologies, Middlesex, UK) per manufacturer’s protocol. Transformed cultures were incubated in Terrific Broth with kanamycin at 37°C overnight (∼16 h). Following this, 10 mL of this culture was added to a 1 L flask containing Terrific Broth together with 20 mL of 20% glucose, 50 mL 1 M KPO_4_, and 50 mg of kanamycin. This culture was grown at 37°C and 250 rpm until the optical density was 0.8-1.0 (∼4-5 h). The incubation temperature was then reduced to 18°C and rotation reduced to 180 rpm for 1 h followed by addition of 50 mL of glycerol and incubation for another 1 h. The culture was induced using 0.1 mM IPTG for 16 h. The cells were pelleted and stored at -80°C until processing and purification.

Pelleted cells were resuspended in Buffer A (100 mM NaCl, 20 mM Tris-HCl at pH 7.5) with EDTA-free protease inhibitor (1 tablet in 50 mL Buffer A per liter of culture). Cells were lysed using an Avestin C5 emulsifier at 15,000 psi with the resuspended cells passing through the emulsifier three times. Cell debris was removed by centrifugation at 14,000 g and 4°C for 15 min. Membranes were collected by ultracentrifugation at 40,000 rpm and 4°C for 45 min. Membranes were homogenized with 45 mL of Buffer A, 5 mL of 10% glycerol, and EDTA-free protease inhibitor, then stored at -80°C until purification. Homogenized membranes were thawed on ice and DDM was added to a final concentration of 1% (v/v). The mixture was rocked at 4°C for 2 h, then ultracentrifuged for 30 min at 40,000 g and 4°C. Amylose resin (New England Biolabs), incubated with Buffer A and 0.05% DDM, was added to the supernatant and rocked at 4°C for 2 h. After rocking, the mixture was centrifuged for 5 min at 500 g and 4°C and the supernatant was removed. Amylose resin beads were placed on a column and the column washed with 20 bed volumes of wash buffer (Buffer A with 0.05% DDM, 1 mM EDTA, 0.5 mM TCEP). Protein was eluted with 5 bed volumes of elution buffer (Buffer A with 0.05% DDM, 0.05 mM TCEP, and 40 mM maltose). Maltose binding protein was removed via digestion overnight with HRV-3C protease (Thermo Fisher, Waltham, MA) (10 units per mg ELIC) at 4°C and purified over a Sephadex 200 Increase 10/300 (GE Healthcare Life Sciences, Pittsburgh, PA) size exclusion column in Buffer A with 0.05% DDM.

### Incorporation of ELIC into Liposomes

A mixture containing 2:1:1 POPC:POPE:POPG in chloroform was placed into a clean glass vial using a glass syringe at room temperature and dried under a gentle stream of N_2_ followed by further drying overnight (∼16 h) in a vacuum desiccator to remove any residual solvent. The resultant thin lipid film was rehydrated with Buffer B (150 mM NaCl, 25 mM MOPS, 0.5 mM BaCl_2_, pH 7.0 adjusted by addition of NaOH) to a final density of 5 mg/mL and vortexed for 5 min. The lipid solution was then subjected to 5 freeze-thaw cycles, alternating between liquid nitrogen and a 40°C heat bath, followed by bath sonication for 1 min. The liposomes were then stored overnight at 4°C. The liposomes were destabilized by adding 0.2% (v/v) DDM and rotating at room temperature for 1 h. Purified ELIC protein was then added to the destabilized liposomes in a 1:25 (protein:lipid) mass ratio and incubated at room temperature for 30 min. Following incubation, DDM was removed from the liposomes by slow addition of BioBeads. Specifically, aliquots of BioBeads (30, 30, and 50 mg) were added to the ELIC proteoliposomes together with 1 mL of Buffer B and rotated at room temperature. Following this, 100 mg of BioBeads were added to the diluted ELIC proteoliposomes and rotated at 4°C overnight (∼16 h). Another 100 mg of BioBeads was added the following morning and rotated at room temperature for 2 h. The ELIC proteoliposomes were separated from the BioBeads and harvested by ultracentrifugation at 45,000 rpm for 1 h at 4°C. The resultant pellet was resuspended in Buffer B (to achieve a lipid concentration of 25-50 mg/ml), divided into 10 *µ*L aliquots, and stored at -80°C until use.

### Single Channel Electrophysiology

An aliquot of ELIC proteoliposomes was removed from -80°C and allowed to come to room temperature. The proteoliposomes were resuspended by gentle trituration and then transferred to a glass cover slip (22 mm×50 mm, Fisher Scientific, Pittsburgh, PA) on top of a bed of calcium sulfate dessicant (Fisher Scientific, Fair Lawn, NJ). The proteoliposomes were placed into a vacuum dessicator for 1 h and dried into a thin film. After removal from the vacuum dessicator, 150 *µ*L of Buffer B was added to the thin film and the proteoliposomes allowed to rehydrate overnight at room temperature. For the giant liposome patch-clamp recordings, a bath solution was prepared with 10 mM cysteamine in Buffer B. Importantly, because cysteamine can oxidize and dimerize (Gonzalez-Gutierrez et al., 2012), a fresh bottle of cysteamine was used for these experiments. The fresh bottle was made into a 5 M solution and aliquots were made and stored at -80°C until use on the day of the experiment. The ELIC proteoliposomes were applied to the bath and allowed to settle for 45 min followed by multiple bath washings with Buffer B. Borosilicate glass capillaries (1B150F-4, ID 0.84 mm, OD 1.5 mm, World Precision Instruments, Sarasota, FL) were pulled using a P-2000 micropipette puller (Sutter Instrument Co., Novato, CA) to a resistance of 7-9 MΩ and Buffer B was added to the pipette. Single channel currents were obtained in an excised patch configuration after formation of a multi-GΩ seal using an AxoPatch 200B (Molecular Devices, San Jose, CA) with the internal low-pass Bessel filter at 5 kHz. The traces were digitized using a Digidata 1440a (Molecular Devices, San Jose, CA) at a sampling rate of 20 kHz and stored using Clampex (version 10.7.0.3). The single channel currents were calculated by fitting an all-points histogram of channel openings to a sum of two Gaussian curves using the ClampFit (version 10.7.0.3) software.

### Modeling and Equilibration of the ELIC5 Open State for Molecular Dynamics Simulation

The open-channel structure of ELIC5 (Petroff II et al., 2022) in an spNW15 nanodisc (PDB 8TWV) (Dalal et al., 2024) was used to model the open-channel state of ELIC with all five mutations (P254G, Y261V, S300C, F319G, F320I) retained. Propylamine was placed in all five orthosteric sites with parameters for propylamine as previously described (Dalal et al., 2024) generated by the CGenFF server (Vanommeslaeghe and MacKerell, Jr., 2012; Vanommeslaeghe et al., 2012). Penalty scores provided by CGenFF server were sufficiently low and did not require further optimization. The N-terminus (P11) of ELIC5 was acetylated and the C-terminus was left uncapped. The local pKa of ionizable groups in ELIC side chains was determined with PROPKA3 (Olsson et al., 2011); this resulted in no protonation/deprotonation of any side chains in the protein with the system pH of 7.0. ELIC5 was then placed in a POPC membrane (539 POPC lipids in the upper leaflet, 545 POPC lipids in the lower leaflet) using CHARMM-GUI (Jo et al., 2008, 2007) and solvated in an aqueous solution consisting of TIP3 waters with 150 mM NaCl. The PPM server (Lomize et al., 2022) was used to orient the protein in the membrane such that the ion conduction pore was along the membrane normal. The initial system dimensions were 210×211×164 Å^3^ with 616,368 atoms.

The ELIC5 system was equilibrated for 50 ns. For systems in which the cryo-EM structure of ELIC5 was under investigation (computational electrophysiology and adaptive biasing force calculations) the backbone atoms of ELIC5 were harmonically restrained (*k* = 5 kcal*/*mol/Å^2^ to their initial positions for 50 ns. For systems in which the dynamics of unrestrained dynamics of ELIC5 were studied, the backbone atoms of ELIC5 were harmonically restrained to their initial coordinates (*k* = 5 kcal*/*mol*/*Å^2^) for 5 ns. The restraints were slowly released over 20 ns with reduction in the harmonic force constant by 0.5 kcal/mol/Å^2^ every 2 ns followed by unrestrained simulation for 25 ns. The resulting system was then subjected to 1 *µ*s of unrestrained molecular dynamics (MD) simulation. This simulation length allowed assessment of appropriate equilibration times for this system as well as determination of the stability for this potentially open structure. Each propylamine was kept in its binding site throughout the full simulation via a weak (*k* = 0.25 kcal*/*mol/Å^2^) bond between the terminal quaternary ammonium of propylamine and the *δ*-carbon of residues E77 and E131. Simulations were carried out in an NPT ensemble. Pressure was maintained at 1 atm using the Nośe-Hoover Langevin piston method (Feller et al., 1995; Martyna et al., 1994) with a piston period of 200 fs and piston decay of 100 fs. Temperature was maintained at 310 K with Langevin dynamics and a damping coefficient of 1 ps^−1^. A 2 fs timestep was used for integration. Short-range non-bonded interactions were cutoff after 12.0Å with a switching function applied after 10.0 Å. Long-range electrostatics were handled using the particle mesh Ewald sums method (Darden et al., 1993). MD simulation was carried out with NAMD 2.14 (Phillips et al., 2020). The CHARMM36 (Klauda et al., 2010) parameter set was used for lipids and ions with CHARMM36m (Huang et al., 2017) and cation-*π* corrections (Khan et al., 2019) being applied to protein segments.

### Modeling of the α7 nAChR and α1 GlyR Open-Channel Structures

In contrast to prokaryotic pLGICs, which have a significantly shortened ICD, eukaryotic pLGICs have an elongated ICD inserted between the M3 and M4 helices that contains a large disordered region. While regions of the ICD have been shown to impact single channel conductance in cationic pLGICs, such as nAChR and 5-HT_A_R (Castelán et al., 2006; Gharpure et al., 2019; Hales et al., 2006; Kelley et al., 2003), the residues implicated are located within the structured regions (*i.e.*, the MA and MX helices) of the ICD which have been resolved. In contrast, point mutations (Ivica et al., 2021), ICD chimeras (Moroni et al., 2011), and ICD deletion (Lara et al., 2019) do not affect single channel conductance of GlyR. Therefore, we did not model the missing disordered regions for either human *α*7 nAChR or zebrafish *α*1 GlyR. For the human *α*7 nAChR model, disulfide bonds were made between C127-C141 and C189-C190. The N-terminus of *α*7 nAChR was resolved and was left charged, but the C-terminus was not and F478 was acetylated. In treating the missing loop of the disordered region, L320 was acetylated with I413 made neutral. For the zebrafish *α*1 GlyR, D100 and D113 were protonated based on PROPKA3 (Olsson et al., 2011) calculations. A disulfide bond was added between C214-C225. Neither the N-nor C-terminus of *α*1 GlyR were resolved, so A25 and V364 were acetylated. In treating the missing loops of the disordered region, both A326 and K329 were acetylated. Each channel was placed into a POPC membrane with CHARMM-GUI (Jo et al., 2008, 2007) and solvated to 150 mM NaCl as above. Harmonic position restraints (k = 5 kcal/mol/Å^2^) were placed on all backbone atoms during initial equilibration of the system as well as throughout all computational electrophysiology calculations for these two models. Initial equilibration of the system proceeded over 50 ns and the last frame of the initial equilibration was used in all computational electrophysiology studies. The temperature used for these simulations was 300 K. All other simulation parameters are as above.

### Computational Electrophysiology

In order to test the conductive properties of the ELIC5 structures, we used the fixed electric field approach for computational electrophysiology (Gumbart et al., 2012; Roux, 2008). While other methods to test the conductive properties of ion channels via simulation have been described (*i.e.*, the double-membrane method (Kutzner et al., 2011) and interpolation from free energy surfaces via the electrodiffusion model (Pohorille and Wilson, 2021)), the fixed electric field method has been used extensively for pLGICs (Avstrikova et al., 2025; Basak et al., 2018; Cerdan and Cecchini, 2020; Cerdan et al., 2018, 2022; Dämgen and Biggin, 2020; Zhuang et al., 2022) and direct comparison with the double-membrane method (Zhuang et al., 2022) demonstrates equivalent results. Computational electrophysiology simulations were performed under an NPT ensemble with the same parameters as above using NAMD 2.14 for the cryo-EM ELIC5 structure and NAMD 3.0 for all other structures. The cryo-EM ELIC5 structure was taken as the last frame of the initial equilibration procedure outlined above. The other ELIC5 structures subjected to the fixed electric field were extracted at 555 ns and 700.8 ns of the unrestrained trajectory. The nAChR *α*7 and *α*1 GlyR structures subjected to the fixed electric field were taken as the last frame of equilibration as outlined above. The fixed electric field was applied in the direction of the membrane normal (*z*-direction) and the strength of the electric field (E*_z_*) was modified to produce transmembrane potentials of either -250 mV, -100 mV, 0 mV, +100 mV, and +250 mV (given that ΔV_m_ = E*_z_*·L*_z_*, where L*_z_* is the length of the simulation box (Gumbart et al., 2012)). In all computational electrophysiology simulations, the protein backbone atoms were subjected to a heavy harmonic restraint (k = 5.0 kcal/mol/Å^2^) throughout the simulation to prevent distortion due to the high transmembrane potentials applied. Each structure was simulated at the indicated transmembrane potential for 150 ns with at least three replicates at each transmembrane potential (*i.e.*, each transmembrane potential examined has at least 450 ns of simulation data). Ion permeation events were counted using an in-house Tcl script (provided in the data repository mentioned under “Data Availability”) and current was estimated as the number of permeation events per total simulation time. Raw data (ion permeation events) was analyzed with an in-house Python script (provided in the data repository mentioned under “Data Availability”). The conductance at positive and negative potentials was measured by estimating the line of best fit through all currents measured for positive (0 to +250 mV) or negative (-250 to 0 mV) transmembrane potentials and taking the slope as the single channel conductance for that structure.

### Adaptive Biasing Force Calculations

To quantify the energetics of ion conduction in the cryo-EM structure of ELIC5 and identify locations of energetic barriers, adaptive biasing force (ABF) calculations (Darve et al., 2008; Hénin et al., 2010) of sodium ion translocation through the channel were performed along two dimensions, namely along the ion conduction axis (*z*-axis) and perpendicular to the ion conduction axis (*xy*-plane). The dimensions were set relative to the I252 C*α* center-of-geometry. Therefore, the center of each window’s *xy* disc was set to the *x*-and *y*-coordinates of the I252 C*α* geometry. The cryo-EM ELIC5 structure being used in the calculation was equilibrated as above and was taken as the last frame of the initial equilibration. The entire length of the channel was divided into windows measuring 10Å in the *z*-direction and 10Å radius in the *xy*-plane with the center of the *xy*-disc centered at the middle of the ion conduction pore. For windows in the TMD portion of the pore in which the radius was much less than 10 Å, the *xy*-plane used for ABF was reduced to 8Å. Within each window, the sodium ion was subjected to flat-bottom harmonic constraints (k = 10 kcal/mol/Å^2^) that only acted if the ion moved outside the boundaries of designated ABF window. The average force acting on the ion was accumulated in bins measuring 0.1×0.1Å^2^ and the adaptive biasing force was applied after 250 samples in each bin. Each window was simulated until convergence of the 2D potential of mean force (PMF) with at least 30 ns of sampling per window. Convergence was obtained by maintaining the root mean-squared deviation (RMSD) of the 2D PMF below 0.1 kcal/mol for at least 5 ns with plateau of both the sampling entropy and sampling heterogeneity (Hénin, 2021). ABF calculations were performed with the Colvars (Fiorin et al., 2013) package and NAMD 2.14 under an NPT ensemble (pressure and temperature control parameters are the same as above) using a 2 fs timestep. A total of 21 windows and 1.025 *µ*s of simulation time were needed for convergence of all windows. The minimum free energy pathway for the apical and lateral pathways were calculated using the string method (E et al., 2007) using fixed endpoints in the extra- and intracellular solution.

## Results

### The cryo-EM ELIC5 structure is stable on the microsecond timescale

ELIC5 is a penta-mutant (P254G/C300S/V261Y/G319F/I320F) construct which does not desensitize, and deactivates on a very slow timescale (Petroff II et al., 2022), suggesting that structures solved in the presence of agonist should represent an open-channel conformation. Several examples of purported open-channel structures of pLGICs, however, have spontaneously and rapidly collapsed to a dehydrated, non-conducting conformation when interrogated with MD simulation (Cerdan et al., 2018; Cheng and Coalson, 2012; Li et al., 2023; Nury et al., 2010). Several independent groups have described use of complex, symmetry-based restraints to bias these channels to keep the pore hydrated (Cerdan et al., 2018; Dämgen and Biggin, 2020; Li et al., 2023). To test the stability of the ELIC5 open-channel structure, we performed equilibrium MD simulation for over 1 *µ*s and measured the root mean-squared deviation (RMSD), hydration number, and ion permeation events (Fig. 1).

**Figure 1.**
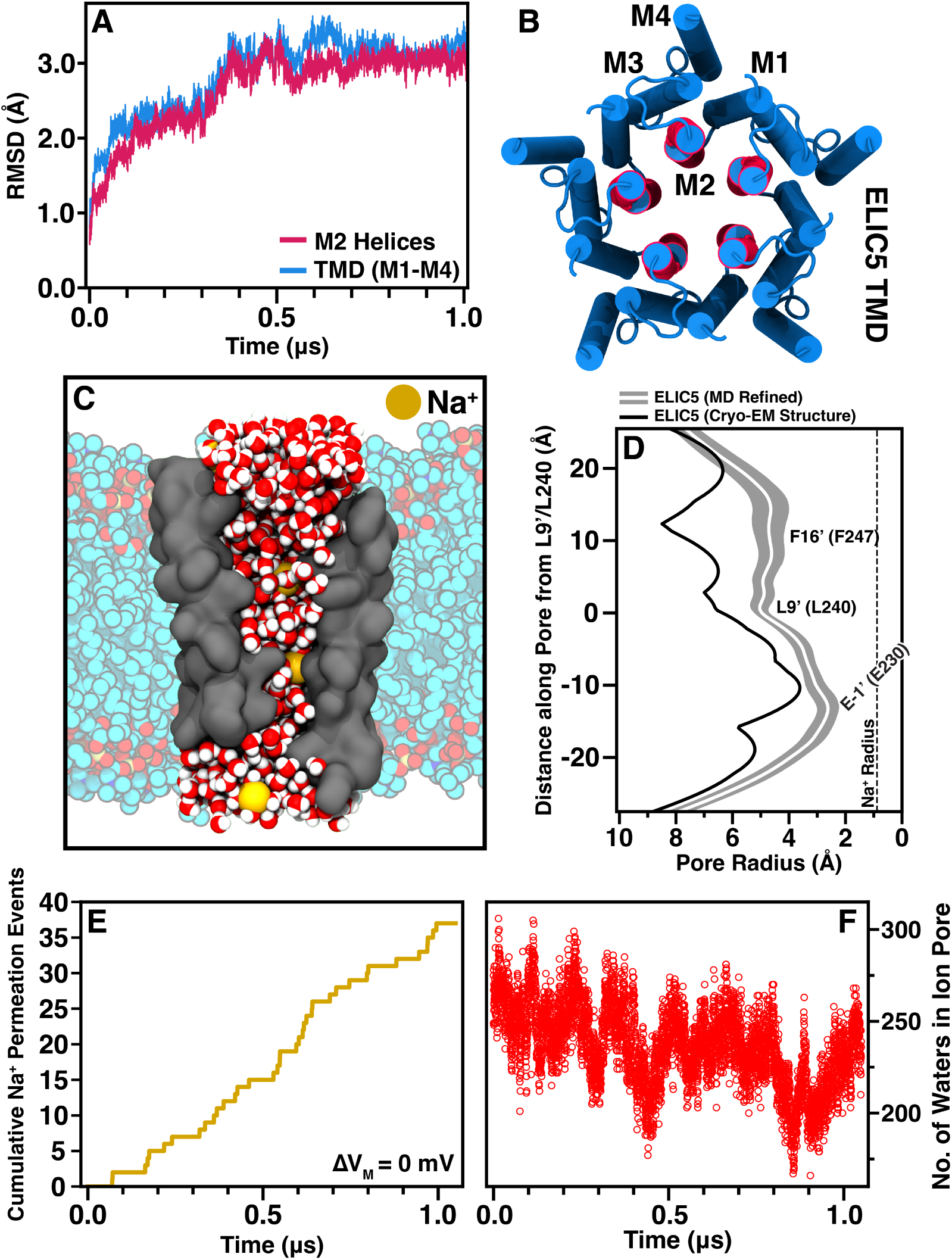
Cryo-EM ELIC5 is stable on the microsecond timescale. (A) Plot of the RMSD of the TMD (blue) and M2 (red) helices during unrestrained MD with a molecular image (B) displaying the protein segments used for the measurement. (C) A snapshot image of ELIC5 from the unrestrained simulation showing a hydrated pore (red/white water molecules) and Na^+^ ions traversing the pore (gold). Only the M2 helices of ELIC5 (gray surface) are shown for clarity. POPC lipids are shown as multi-colored models in the background. (D) The ion conduction pore radius of ELIC5 over the last 500 ns of unrestrained simulation as measured with HOLE is shown (mean pore radius- white line, standard deviation-gray area) along with the starting ion conduction pore radius (black) measured from the cryo-EM structure (PDB 8TWV). The position of the hydrophobic gate residues (L9’ and F16’) as well as the selectivity filter (E-1’) are marked. The ionic radius of an unsolvated Na^+^ ion is shown as a dashed line. The cumulative number of Na^+^ ions traversing the ion conduction pore (E) as well as the total number of waters in the ion conduction pore (F) are shown for the duration of the unrestrained simulation. No transmembrane potential was applied for these unrestrained simulations.

It is not well-established how long pLGICs require for equilibration from their static structures, but studies using long timescale simulations (Avstrikova et al., 2025; Hénault et al., 2019) have demonstrated ongoing structural rearrangements at 300-400 ns. Our simulations show that equilibration of the channel is reached only after 500-600 ns with a stable plateau in RMSD reached after this time point for all domains (Fig. 1A). Examining the pore radius over the last 500 ns of the simulation trajectory (*i.e.*, after the structure had equilibrated), shows that there has been reduction in radius along the whole TMD (Fig. 1D). The narrowest point is still located at E230 (a conserved ring of glutamates at the -1’ position of the M2 helix) with an average radius of 2.4± 0.3Å. This makes the channel still wide enough to conduct Na^+^ ions even after equilibration. The hydrophobic residues at 9’ and 16’, L240 and F247, respectively, are the activation gates of ELIC based on the resting structure. The pore diameter at these residues is wide open in the ELIC5 open-channel structure, and there is a slight narrowing after equilibration of the simulation. Importantly, the ion channel pore remains hydrated over the length of the simulation and Na^+^ ions are seen to traverse the channel throughout the equilibrium simulation as well, even in the absence of an applied transmembrane potential (Fig. 1E,F). Taken together, these data show the ELIC5 construct used in these simulations is a stable, conducting conformation. While these are necessary conditions for the physiologic open-channel state, they are not sufficient. Further evidence is needed to demonstrate that sodium conduction through the channel matches the experimental value.

### Sodium conductance in cryo-EM ELIC5 does not match functional data

To test whether the ELIC5 open-channel structure from cryo-EM (hereon termed ”cryo-EM ELIC5”) represents the physiologic open-channel state, we examined sodium conduction with computational electrophysiology. In this approach, a fixed electric field is applied across the length of the simulation box, resulting in a fixed transmembrane electric potential (ΔV_m_, Fig. 2A). Due to the timescale limitations of MD studies, relatively high ΔV_m_ are applied (ranging from -250 to 250 mV in this study) in order to observe sufficient ion crossings. To prevent distortion of the channel structure, harmonic restraints are placed on the backbone atoms. Upon applying a range of positive and negative ΔV_m_ to cryo-EM ELIC5, a large outward rectification is observed (Fig. 2B) with an outward conductance (*γ_o_*) of 85.5 ± 4.9 pS, an inward conductance (*γ_i_*) of 21.0 ± 3.8 pS, and a rectification ratio (*γ_o_*/*γ_i_*) of 4.07 (Table 2). This behavior was reproducible over multiple simulations (n=3 independent simulations for each potential). While *γ_o_* agrees with experimental conductance measurements, *γ_i_* is much lower than expected. Wild-type ELIC is not known to rectify (Zimmermann and Dutzler, 2011) and its Ohmic behavior is redemonstrated here by excised patch-clamp recordings of ELIC from giant liposomes (Fig. 2C,F).

**Figure 2.**
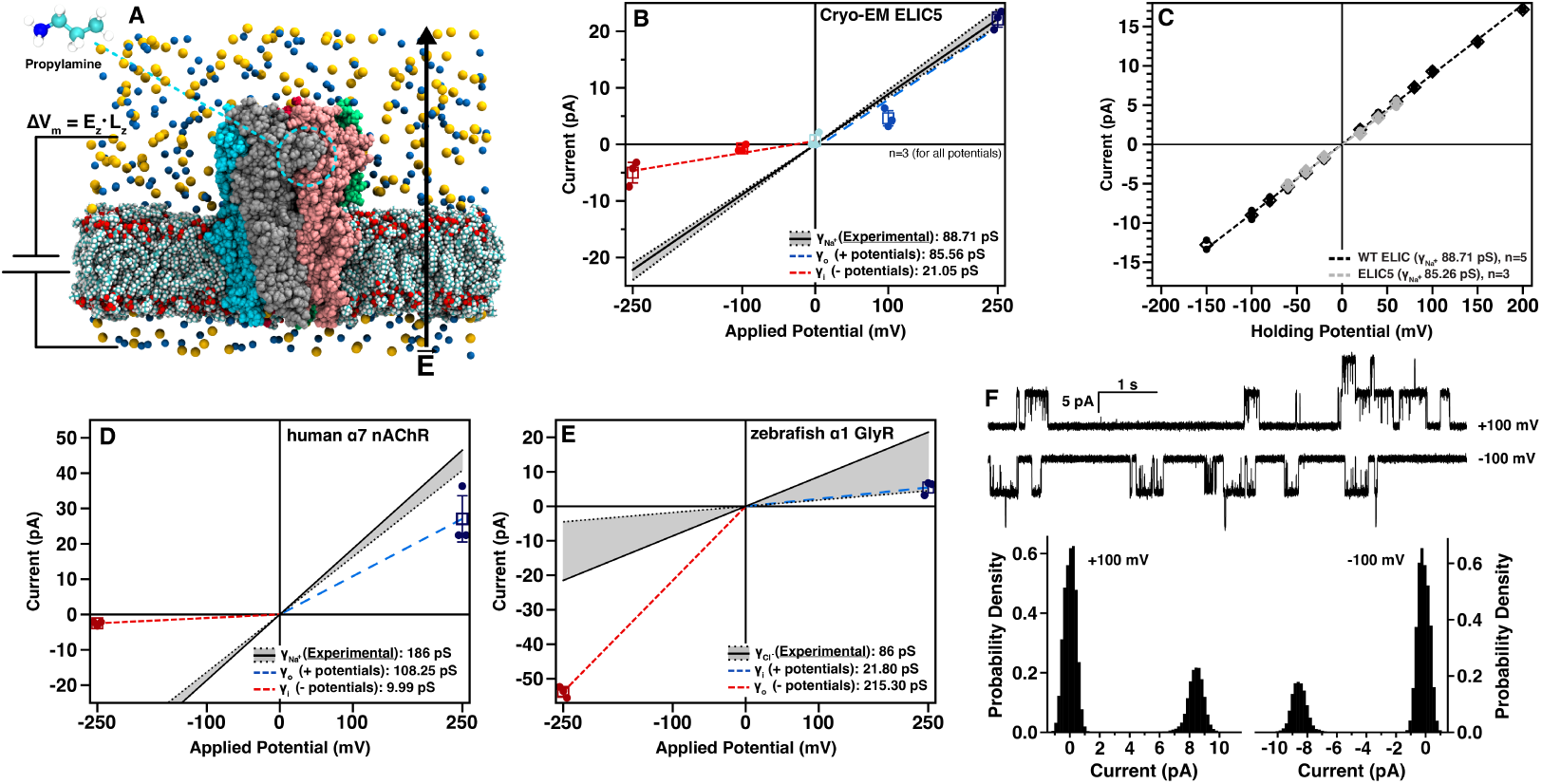
The cyo-EM structures of pLGICs display significant outward rectification. (A) A molecular schematic for the applied transmembrane potential simulations. The electric field is applied only in the direction of the membrane normal. ELIC5 is shown colored by subunit with sodium ions in yellow, chloride ions in blue, and POPC as multicolored van der Waals surfaces. Propylamine is placed in all five agonist binding sites. (B) Measured currents for each transmembrane potential of cryo-EM ELIC5. Independent replicates are shown as dots, mean as a box, standard deviation as a crossbar, and conductance for positive and negative potentials as blue and red dashed lines, respectively. The experimental single-channel Na^+^ conductance (*γ*_Na_) is shown as a solid black line (standard deviation in gray). (C) Current-voltage relationship for WT ELIC (black, n=5) and ELIC5 (gray, n=3) from excised patch-clamp recordings of giant liposomes. Replicates are shown as dots, the mean as a diamond, and the standard deviation as a crossbar. The dashed lines represent the line of best fit through the measured mean currents and is taken as *γ*_Na_. The results of computational electrophysiology measurements for (D) human *α*7 nAChR (PDB 7KOX) and (E) zebrafish *α*1 GlyR (PDB 3JAE) are shown with the same color scheme as in (B). (F) Representative single-channel currents of WT ELIC (top), and histograms of the currents from these traces (bottom).

**Table 2:** Single channel conductance and rectification ratio for ELIC and other pLGICs from computational electrophysiology studies. *γ_o_* is estimated using a linear fit of all data points from 0, +100, and +250 mV; *γ_i_* is estimated using a linear fit of all data points from 0, -100, and -250 mV. The error in the plot represents the standard error from the linear fitting.

| Structure | PDB | $\gamma_o$ (pS) | $\gamma_i$ (pS) | Rectification Ratio |
| --- | --- | --- | --- | --- |
| ELIC5 (Cryo-EM) | 8TWV | $85.5 \pm 4.9$ | $21.0 \pm 3.8$ | 4.07 |
| ELIC5 (MD Relaxed 1) | – | $160.0 \pm 8.6$ | $91.4 \pm 5.0$ | 1.75 |
| ELIC5 (MD Relaxed 2) | – | $172.6 \pm 6.6$ | $104.8 \pm 7.4$ | 1.65 |
| human $\alpha 7$ nAChR | 7KOX | $108.2 \pm 15.1$ | $9.9 \pm 1.1$ | 10.84 |
| zebrafish $\alpha 1$ GlyR | 3JAE | $21.8 \pm 3.0$ | $215.3 \pm 3.6$ | 0.10 |

Likewise, ELIC5 has the same *γ*_Na_ as WT ELIC (88.7 ± 1.6 pS) and did not display any rectification (Fig. 2C, Table 3). Therefore, ELIC5 is suitable for examining the sodium conduction properties of WT ELIC, but computational electrophysiology of cryo-EM ELIC5 shows an outward rectification that does not agree with experiment.

**Table 3:** Single channel conductance and rectification ratio for ELIC constructs. For each patch, outward (*γ_o_*) and inward (*γ_i_*) conductances were estimated by taking the slope of the linear fit for all currents at either positive or negative ΔV_m_, respectively. The mean was calculated as an average across all patches and the error presented is the standard deviation. The rectification ratio represents *γ_o_/γ_i_*. Statistical significance between *γ_o_* and *γ_i_* in the same construct was determined using a paired student’s t-test. Statistical significance between *γ_o_* and *γ_i_* is denoted by (*) in the “Rectification Ratio” column. Statistical significance between mutants was determined separately for *γ_o_* and *γ_i_* using one-way ANOVA. One-way ANOVA F-statistic for *γ_o_* (13.4806*>*2.57 for *α*=0.05) and *γ_i_* (173.5423*>*2.57) indicated significance, prompting Dunnett’s multiple comparisons test (*α*=0.05) to determine which mutants were statistically significant when compared to WT. Statistical significance from WT is indicated by (†).

| | $\gamma_o$ (pS) | $\gamma_i$ (pS) | Rectification Ratio | No. of Patches |
| --- | --- | --- | --- | --- |
| ELIC | $89.76 \pm 3.84$ | $84.99 \pm 2.60$ | 1.06 (p=0.09) | 5 |
| ELIC5 | $94.93 \pm 4.13$ | $91.38 \pm 2.92$ | 1.04 (p=0.07) | 3 |
| R65A | $86.18 \pm 4.96$ | $102.82 \pm 2.82^\dagger$ | 0.83* (p=0.0007) | 7 |
| R65A/K90A | $89.11 \pm 7.84$ | $125.71 \pm 7.92^\dagger$ | 0.70* (p=0.0002) | 6 |
| E30A/D158A | $70.29 \pm 4.85^\dagger$ | $20.54 \pm 4.43^\dagger$ | 3.42* (p=0.0001) | 6 |
| D113A/D158A | $70.52 \pm 5.46^\dagger$ | $32.04 \pm 11.71^\dagger$ | 2.20* (p=0.0059) | 5 |

To determine if the rectification observed in cryo-EM ELIC5 is unique to ELIC, we applied the same computational electrophysiology approach to the open-channel structures of full-length human *α*7 nAChR (Noviello et al., 2021) and zebrafish *α*1 GlyR (Du et al., 2015). In both cases, a significant outward rectification is observed either with sodium conductance in *α*7 nAChR or chloride conductance in *α*1 GlyR (Fig. 2D,E). The results for human *α*7 nAChR reproduce those obtained with a double-membrane approach Zhuang et al. (2022), while the results for full-length *α*1 GlyR have not been previously published. While Na^+^ has been observed to traverse the 5-HT_A_ open-channel state using computational methods (Basak et al., 2018), this was only at one ΔV_m_ and so the rectification of this construct is unknown. These results demonstrate that the behavior observed in cryo-EM ELIC5 are not anomalous, but a trend observed across cryo-EM open-channel structures of pLGICs.

The discrepancy in the inward sodium conduction of cryo-EM ELIC5 and the electrophysiology data prompted further examination of the conducted Na^+^ ions in the equilibrium MD simulation. Visual inspection revealed that several ions carrying inward current (*i.e.*, Na^+^ ions traversing the extracellular to intracellular direction), were not originating from the extracellular side of the simulation box, but rather had been conducted from the intracellular side immediately prior (Fig. 3A). This result suggested that there was normal ion flow through the TMD, but restricted ion flow between the extracellular solution and the central vestibule in the ECD. Therefore, we performed computational electrophysiology simulations as above with a TMD-only construct of the cryo-EM ELIC5 structure. These simulations eliminated the rectification observed in the full-length construct (Fig. 3B), further suggesting restricted ion flow in the ECD. Although the *γ*_Na_ for the TMD-only construct is much higher than the experimental *γ*_Na_ for ELIC5, it is expected that removing the ECD, which has known ion interaction sites (Zimmermann et al., 2012), would increase the *γ*_Na_ compared to the full length receptor. Indeed, this same phenomenon has been described for GlyR (Cerdan et al., 2022).

**Figure 3.**
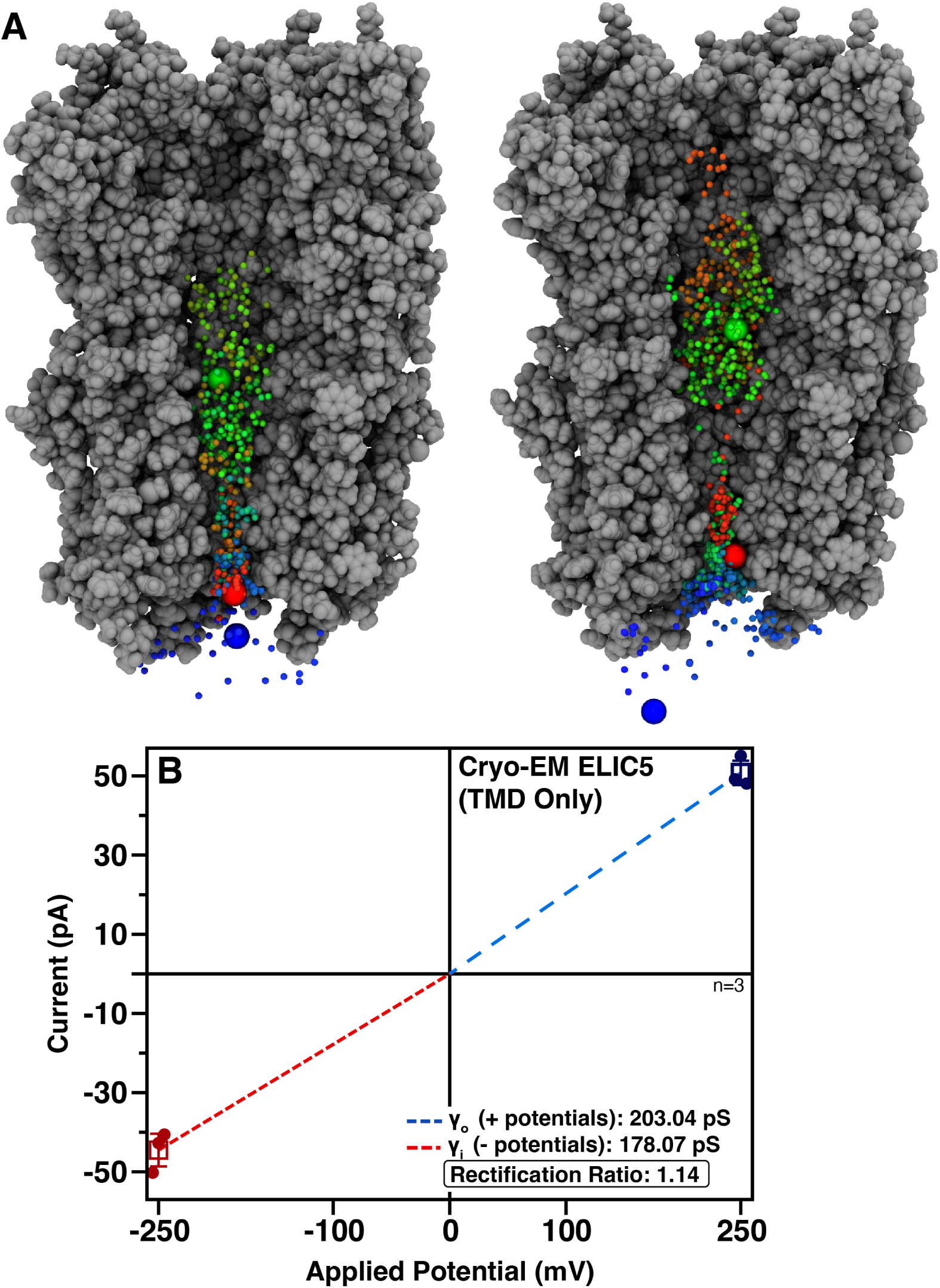
The cryo-EM ELIC5 ECD limits sodium conductance. (A) Two examples of Na^+^ ions traversing the TMD from the intracellular side to the ECD vestibule, but unable to exit the ECD and traversing back across the TMD. Cryo-EM ELIC5 is shown as a gray surface with subunits removed for clarity. A single Na^+^ ion is shown in each panel with its position as a function of simulation time shown as a blue-green-red spectrum. The initial position of the Na^+^ ion on the intracellular side is shown as a large blue sphere. The time point when the Na^+^ ion exits the TMD to the ECD is marked by the large green sphere and the time point when the Na^+^ crosses back across the TMD is marked with a large red sphere. (B) Current-voltage relationship for cryo-EM ELIC5 with the ECD removed (*i.e.*, TMD-only construct) with the same coloring scheme as in Fig. 2B.

### Elucidation of a lateral, intersubunit conduction pathway in ELIC

The computational electrophysiology results suggest restriction of Na^+^ ion flow between the extracellular solution and the central vestibule of ELIC. To locate the origin of the restriction, adaptive biasing force (ABF) calculations were used to determine the potential of mean force (PMF) for Na^+^ conduction through cryo-EM ELIC5 (Fig. 4). For unhindered ion flow, the energy surface for ion translocation should be flat. The PMF shows that there is a relatively flat energy surface for Na^+^ movement through the TMD, in line with the computational electrophysiology results. In the ECD, however, there are significant energy barriers and energy wells preventing ion flow. Along the apical pathway, R65 and K90 form a two-layered energetic barrier to Na^+^ conduction that measures 1.81 kcal/mol at its peak and spans approximately 10 Å at the apical opening. Thus, it appears that ion movement across the apical pathway would be unfavored. Conversely, a lateral fenestration or pathway, located between subunits in the ECD, has a lower energy surface for Na^+^ movement with several deep energy wells, the deepest measuring approximately -7.5 kcal/mol located at the junction of the lateral pathway and central vestibule at residues E30 and D158. Residues E36, D113, D155, D157, and D158 along the lateral pathway form the other energy wells. The deepest energy wells are located where multiple acidic residues come together to coordinate Na^+^ stably. Interestingly, some of these acidic residues, D113 and D158, are known to abolish the inhibitory effects of Ca^2+^ on ELIC activation and crystal structures show multiple bound Ba^2+^ ions at either end of this pathway (Zimmermann et al., 2012). Notably, the lateral pathway elucidated here for ELIC is similar to the one recently described for *α*1 GlyR (Cerdan et al., 2022), but has not been described in other pLGICs to date. The ABF calculations suggest the lateral fenestration would be the main ion conduction pathway in ELIC, which was also proposed for *α*1 GlyR (Cerdan et al., 2022).

**Figure 4.**
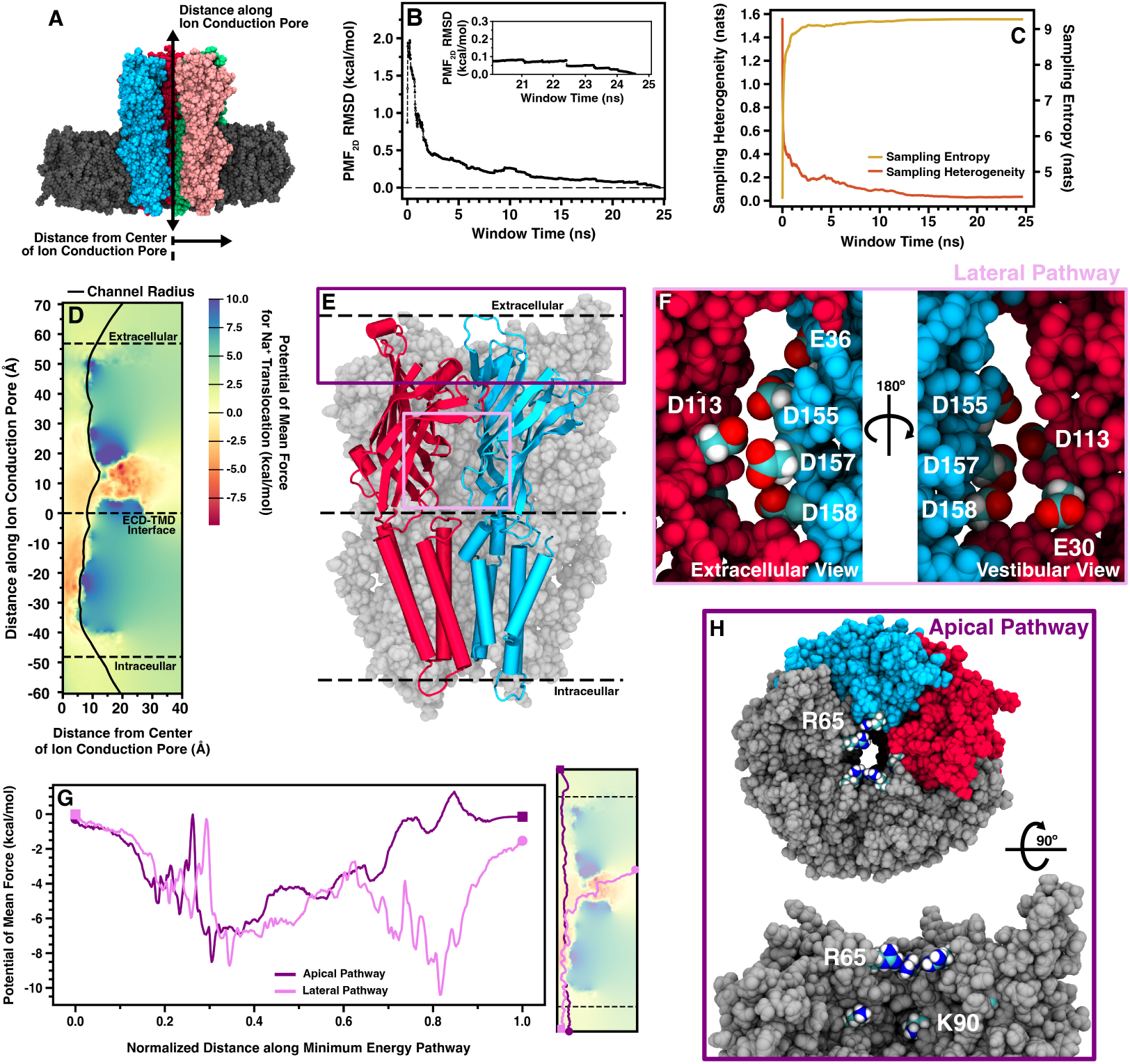
Potential of mean force for Na^+^ in cryo-EM ELIC5. (A) Molecular image showing the two axes used in calculating the 2D PMF for Na^+^ translocation across cryo-EM ELIC5. Representative plots of the 2D PMF RMSD (B), sampling entropy (C, gold trace), and sampling heterogeneity (C, orange trace) used to monitor convergence in each ABF window shown for one window. The inset in (B) shows the 2D PMF RMSD for the last 5 ns in the window. (D) The full 2D PMF for cryo-EM ELIC5 shown with the energy relative to bulk solution. The radius of the entire channel as calculated by HOLE is shown as a solid black line. (E) Molecular image of ELIC5 cryo-EM with two subunits highlighted in blue and red. The location of the apical pathway residues (R65 and K90) is shown as a dark purple box, and the lateral pathway as a light purple box. The acidic residues in the lateral pathway are displayed from the extracellular solution (F, left) and the central vestibule (F, right). (G) The minimum energy pathway (right) and energy along the pathway (left) is shown for the apical (dark purple) and lateral (light purple) pathways. As the two paths have different absolute lengths, the independent axis displays the distance along the minimum energy pathway normalized to the total length of the path. (H) Molecular images of the apical pathway highlighting the location of residues R65 and K90.

Given the results from the ABF calculations, we examined the dynamics of the apical and lateral pathways from the MD simulation (Fig. 5). The lateral pathway has a noticeable widening over the last 500 ns of the trajectory. In the cryo-EM structure, the lateral pathway is too narrow to freely conduct Na^+^ ions. By the last 500 ns, however, the lateral pathway widened sufficiently to freely allow for Na^+^ conduction (Fig. 5A). The apical opening at the levels of both R65 and K90 widen as well (Fig. 5B), suggesting that the cryo-EM structure has an overly compact ECD. With the widening of the lateral pathway, there is separation of acidic residues that form the deep energy wells observed in the ABF calculations, which may reduce or eliminate the restricted ion flow. Therefore, two snapshots from the equilibrated trajectory were selected for analysis by computational electrophysiology (Fig. 6). We chose two structures with different width at the apical pathway to examine whether separating the high positive charge density of R65 and K90 would allow for more conduction via this less favored pathway. In these MD-refined structures, there is a striking reduction in the rectification compared with the cryo-EM ELIC5 structure (1.65/1.75 vs 4.07, Table 2), providing better agreement with experimentally measured rectification ratios. Moreover, the sodium conductance on either side of zero more closely match experimental values for WT ELIC and ELIC5.

**Figure 5.**
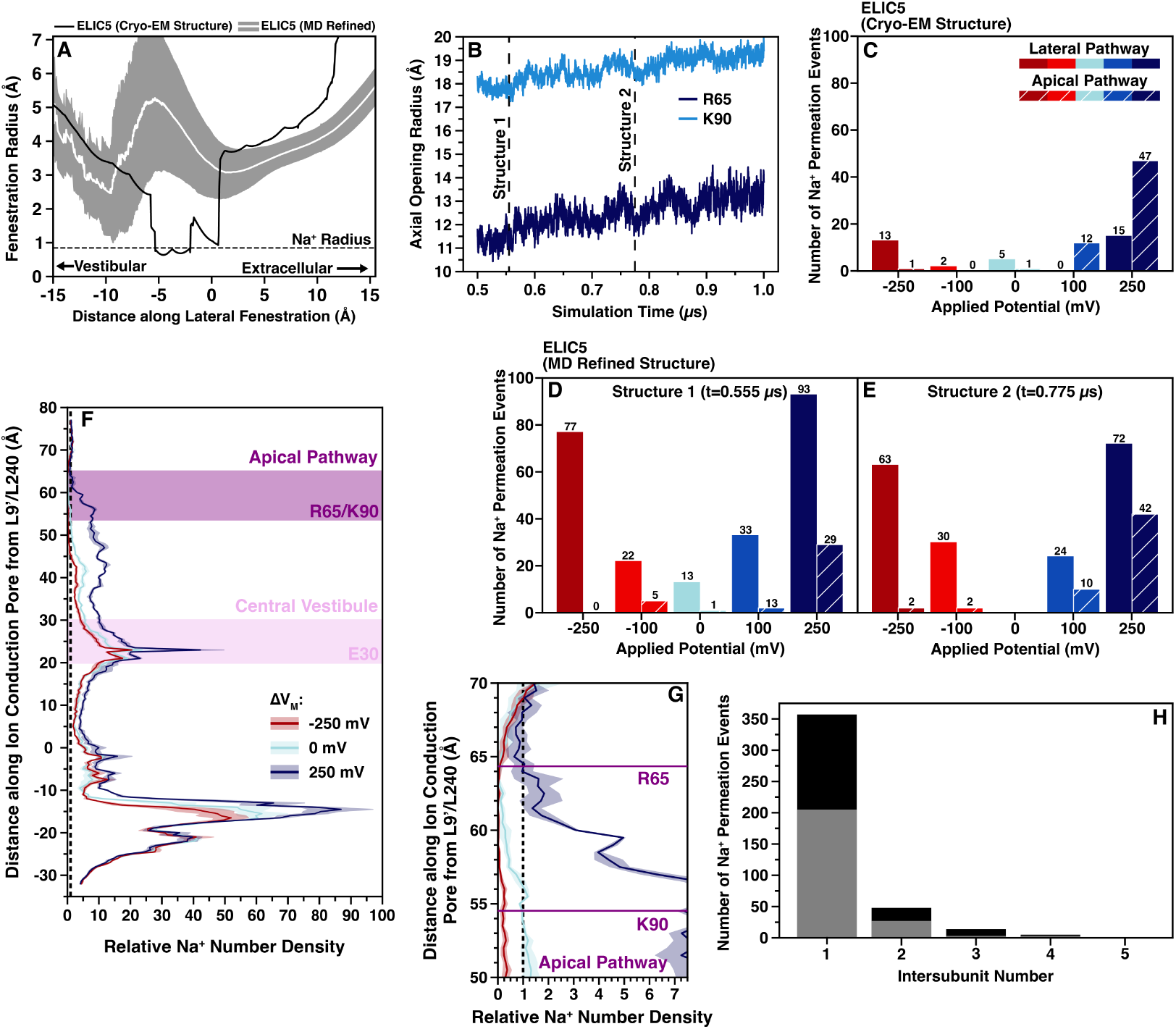
Na^+^ conduction along the lateral and apical pathways. (A) HOLE radius of the lateral fenestration over the last 500 ns of unrestrained simulation of ELIC5 (mean radius- white line, standard deviation-gray area), and the initial HOLE radius of ELIC5 cryo-EM (PDB 8TWV). The radius of an unsolvated Na^+^ ion is shown as a horizontal dashed line for reference. (B) The radius of the residues at the two levels of the apical pathway energetic barrier, R65 (dark blue) and K90 (light blue) over the last 500 ns of unrestrained simulation. The time points used for the MD refined structures in the computational electrophysiology simulations (Structure 1 and Structure 2) are noted as vertical dashed lines. Plots of the number of conducted Na^+^ ions traversing the lateral (solid bars) or apical (hashed bars) pathways is shown for each applied transmembrane potential from computational electrophysiology simulations of cryo-EM ELIC5 (C), ELIC5 MD refined Structure 1 (D), and ELIC5 MD refined Structure 2 (E). (F) Plot of the number density of Na^+^ ions along the channel is shown for cryo-EM ELIC5 at -250 mV (red), 0 mV (light blue), and +250 mV (blue) with the mean number density (n=3) shown as the solid trace and the standard deviation shown as the shaded area. Number density is shown relative to bulk solution. A close-up of the apical barrier region Na^+^ number density is shown in (G). (H) A stacked bar plot showing the number of conducted Na^+^ ions which cross each lateral fenestration for all applied transmembrane potentials is shown for ELIC5 MD refined structure 1 (gray) and structure 2 (black).

**Figure 6.**
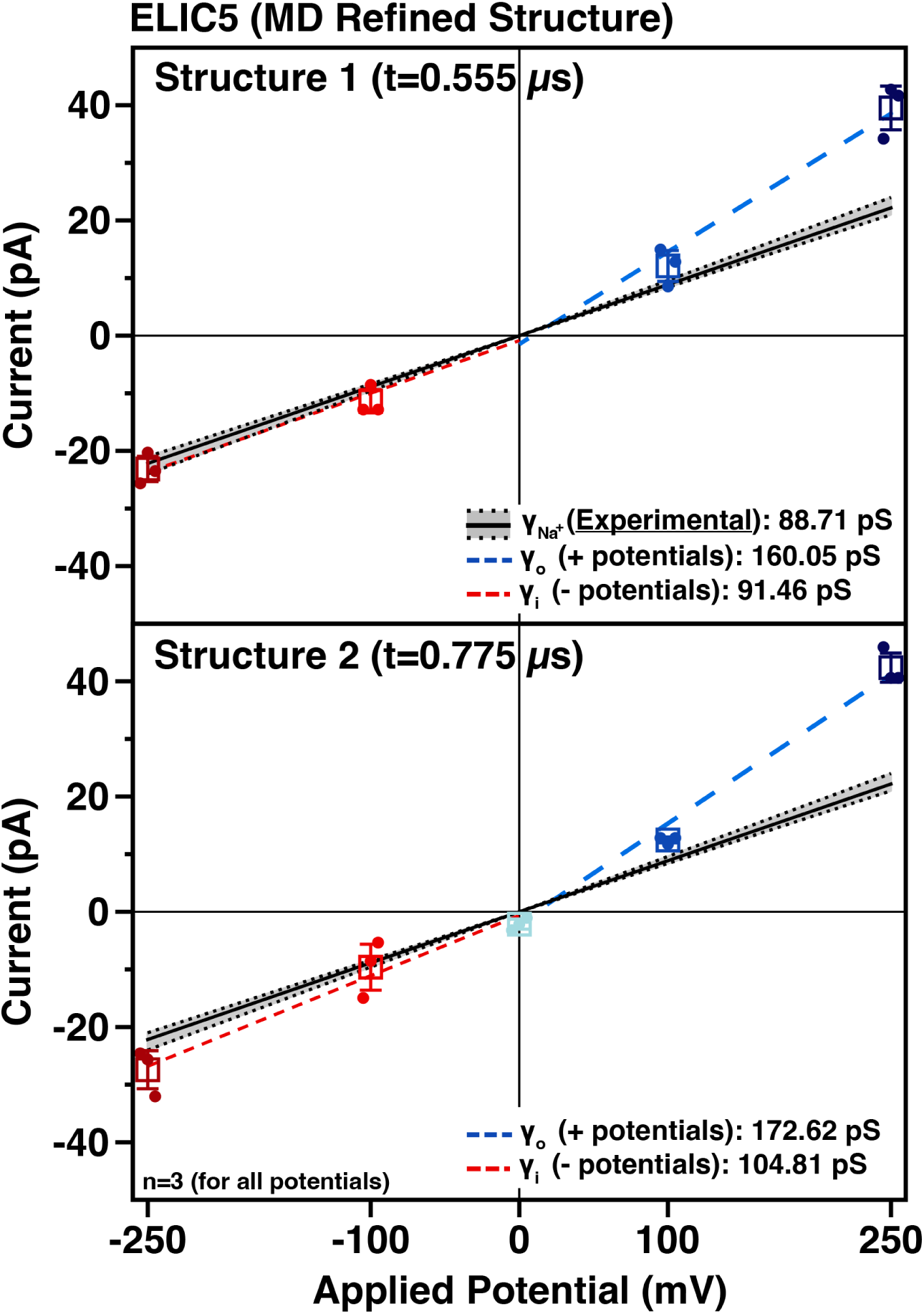
Computational electrophysiology of ELIC5 MD Refined Structures. The current-voltage relationships for the ELIC5 MD refined structures (structure 1, top; structure 2, bottom) are shown with the same coloring scheme as Fig. 2B.

To better understand the relative contributions of the apical and lateral pathways to sodium conduction, the number of conducted ions which passed the apical or lateral pathways were counted (Fig. 5C-E). In cryo-EM ELIC5, the conduction pathway is dependent on the orientation of ΔV_M_ with Na^+^ ions flowing mostly across the apical opening at positive ΔV_M_ versus flowing mostly through the lateral pathway at negative ΔV_M_. A similar trend is observed in the zebrafish *α*1 GlyR structure where apical flux is higher in outward Cl^−^ conductance. In contrast, the MD-refined structures show the majority (∼80%) of conducted Na^+^ ions passing through the lateral fenestration regardless of ΔV_M_. Only at +250 mV is there appreciable conduction via the apical pathway, which may be a result of the large applied potential. Measuring the Na^+^ number density in the ELIC5 MD-refined structures demonstrates that at +250 mV the channel is able to concentrate Na^+^ to a much greater degree in the ECD than at either -250 mV or 0 mV (Fig. 5F), especially around residues in the central vestibule (E30) and the apical pathway barrier (R65/K90) (Fig. 5G). This accumulation of Na^+^ ions in the ECD would likely lead to increased crossing of the apical pathway at positive potentials.

Together, the ABF and computational electrophysiology calculations point to the importance of the lateral pathway in ELIC sodium conductance, especially inward conductance. To validate this, single and double mutants of charged residues identified along the apical (R65, K90) and lateral (E30, D113, D158) pathways were mutated to alanine and their single-channel properties measured (Fig. 7, Table 3). If the computational model of ELIC conduction is accurate, elimination of the basic residues at the apical barrier should increase conductance and elimination of the acidic residues along the lateral pathway should lower conductance. As expected, both the single (R65A) and double (R65A/K90A) apical pathway mutants show increased *γ_i_* and an unchanged *γ_o_*, with the double mutant having a more profound effect on *γ_i_* (Fig. 7). Mutation of the lateral pathway residues, however, decreased both *γ_i_* and *γ_o_*, with a greater effect on the inward conductance. The lateral pathway double mutants, E30A/D158A and D113A/D158A, have an outward rectification measuring 3.42 and 2.20 times that of wild-type, respectively. Therefore, the effects of both the apical and lateral pathway mutants agree with the MD simulation data supporting the key role of the lateral pathway for Na^+^ conduction in ELIC.

**Figure 7.**
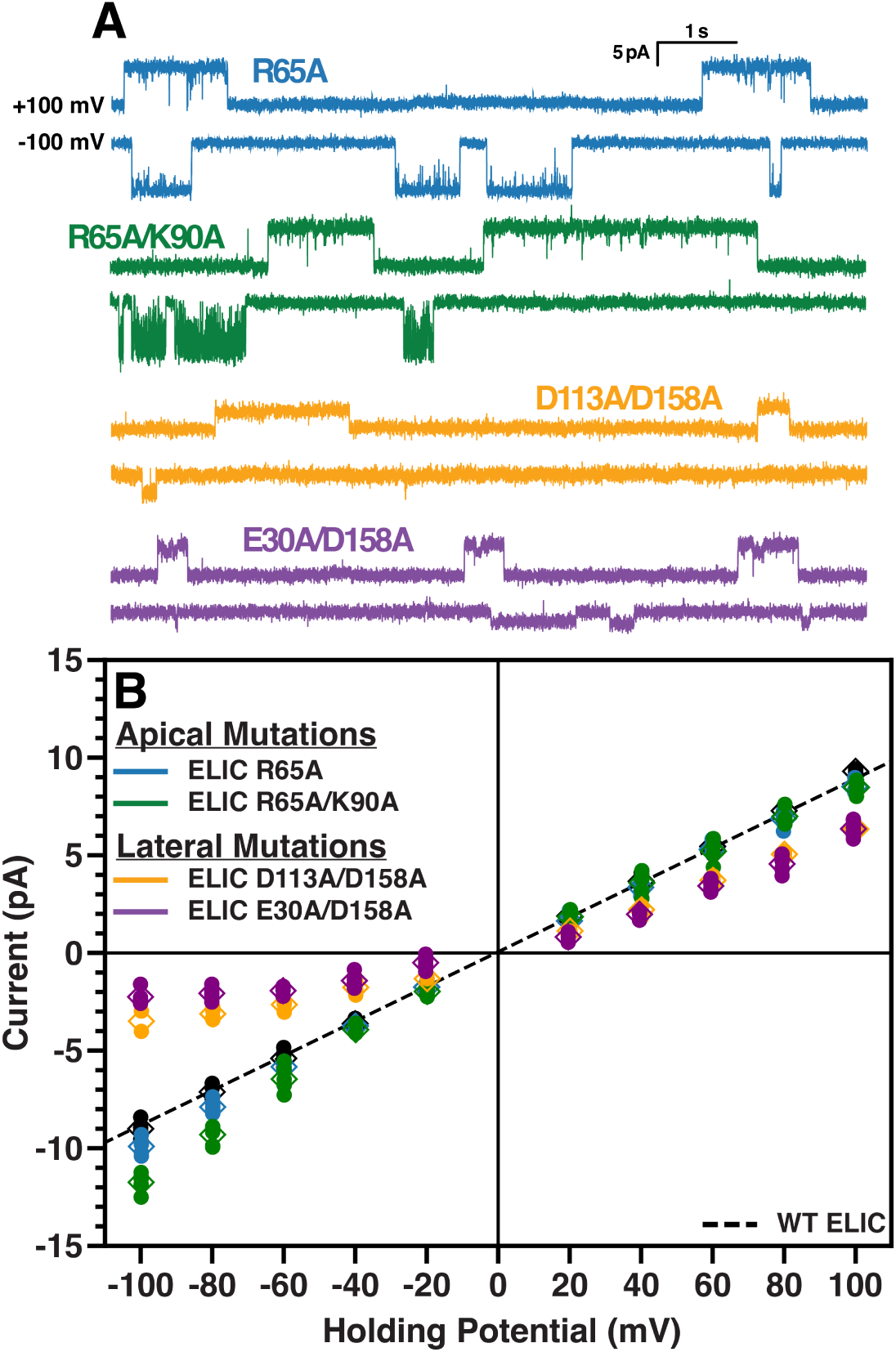
Mutations of acid residues in the lateral pathway reduce single-channel conductance. (A) Representative single-channel currents of ELIC single/double mutants. Data for WT ELIC from Fig. 2C are shown in black. Mutants of the apical barrier residues are shown in blue (R65A) and green (R65A/K90A), and mutants of the lateral fenestration residues are shown in gold (D113A/D158A) and purple (E30A/D158A). (B) Current-voltage relationships for the mutants described in (A) are shown with the same color scheme. The data from individual recordings is shown as dots with the mean as diamonds and the standard deviation as crossbars. Slope conductances were calculated separately for positive and negative potentials for each mutant. The slope conductances are not displayed here for clarity, but are available in Table 3 along with the rectification ratio. The slope conductances are calculated using the dominant single-channel currents from each patch.

## Discussion

Here, we examined the sodium conduction properties of the cryo-EM open-channel structure of ELIC (cryo-EM ELIC5) using a combination of MD simulation and electrophysiology measurements. The results demonstrate that cryo-EM ELIC5 is stable on the microsecond timescale: the pore remains hydrated and able to conduct Na^+^ ions. This is in contrast to several other purported open-channel pLGIC structures which show rapid and irreversible collapse of the TMD upon simulation (Cerdan et al., 2018; Cheng and Coalson, 2012; Li et al., 2023; Nury et al., 2010). However, the computational electrophysiology results indicate that while a channel structure may remain hydrated, this is not sufficient to declare the structure an “open-channel state”. Indeed, it is possible for the structure to demonstrate a much higher conductance than what is known, as in the case of GlyR (Cerdan et al., 2018), or to display significant rectification and a single channel conductance that is lower than what is known, as in the case of ELIC5 (presented here) and *α*7 nAChR (Noviello et al., 2021).

Upon unrestrained MD simulation, a pre-existing, but highly constricted, lateral fenestration spontaneously widens, allowing a significant reduction in outward rectification observed in the cryo-EM ELIC5 structure (PDB 8TWV) as well as better agreement with experimental *γ*_Na_. The importance of the lateral fenestration in *α*1 GlyR (PDB 3JAE) ion conduction has been detailed (Cerdan et al., 2022), but this phenomenon has not been previously described for cation-permeable pLGICs, such as ELIC. This suggests there may be a conserved mechanism for ion conduction across both cation- and anion-permeable pLGICs, although it is not clear that all pLGICs conduct a majority of their permeant ions across a lateral fenestration. In the case of human *α*7 nAChR (PDB 7KOX), there is not an obvious lateral fenestration in the cryo-EM structure and previous examination of ion conduction in the open-channel structure (Noviello et al., 2021) did not mention a lateral ion conduction pathway. This study, however, used harmonic restraints to enforce the geometry of the cryo-EM structure and it is possible that unrestrained simulation may elucidate a cryptic lateral fenestration. Limited examination of ion conduction in mouse 5-HT_A_ (Basak et al., 2018) did not report a lateral ion conduction pathway either. With the recent determination of multiple open-channel structures for pLGICs, there is an opportunity to examine the diversity of ion conduction mechanisms across the pLGIC superfamily.

The outward rectification observed in the cryo-EM ELIC5 model is not unique to this particular pLGIC. When examined using a double-membrane computational electrophysiology approach, human *α*7 nAChR displayed the same outward rectification pattern to an even greater degree (*γ_o_* = 223.0 pS, *γ_i_*= 44.2 pS, rectification ratio= 5.04) (Zhuang et al., 2022). Outward rectification was also observed using a constant electric field approach in this study for *α*7 nAChR (*γ_o_*= 108.25 pS, *γ_i_*= 9.99 pS, rectification ratio= 10.84). Computational electrophysiology data for the purported open-channel structure of full-length zebrafish *α*1 GlyR were not available previously. Previous studies had either used a TMD-only construct (Cerdan et al., 2018) or performed computational electrophysiology on an MD refined structure (Cerdan et al., 2022). Results from this study show the same outward rectification pattern for *α*1 GlyR (*γ_o_*= 21.80 pS, *γ_i_*= 215.30 pS, rectification ratio= 0.10). Recently published structures of GLIC also have demonstrated significant outward rectification, even in asymmetric states (Li et al., 2025). While a purported open-channel structure for 5-HT_A_ exists (Basak et al., 2018), the measured single-channel conductance is low enough (estimated to be ∼400-900 fS) to make the computational approaches used in this study challenging. The results here, and in previous studies, however, suggest there is a common artifact causing compaction of the ECD in cryo-EM open-channel pLGIC structures. This compaction of the ECD leads to a dearth of ions in the central vestibule above the TMD. Our results show that low ion number density in the central vestibule is associated with low inward conductance and outward rectification. Indeed, when the residues that concentrate Na^+^ in the central vestibule (E30, D158) are mutated in ELIC, there is a significant decrease in inward conductance and a resultant outward rectification. Here, MD simulation displays expansion of the lateral fenestration, the main Na^+^ conduction pathway described here, which allows Na^+^ to enter the central vestibule from the extracellular solution and concentrate above the TMD. This removes most of the outward rectification observed in the cryo-EM ELIC5 structure and produces a model more consistent with experimental data; similar results were demonstrated for *α*1 GlyR previously (Cerdan et al., 2022). Therefore, it seems prudent that all open-channel structures of pLGICs should be probed with MD simulations, which allow solvation, ions, and physiologic temperature, when examining structure-function relationships.

## Abbreviations

5-HT_A_: serotonin receptor type 3A
ABF: adaptive biasing force
cryo-EM: cryo-electron microscopy
DeCLIC: *Desulfofustis glycolicus* ligand-gated ion channel
ECD: extracellular domain
ELIC: *Erwinia christanemii* ligand-gated ion channel
GLIC: *Gloeobacter violaceus* ligand-gated ion channel
GluCl: glutamate-gated chloride channel
GlyR: glycine receptor
ICD: intracellular domain
MD: molecular dynamics
nAChR: nicotinic acetylcholine receptor
pLGIC: pentameric ligand-gated ion channel
PMF: potential of mean force
POPC: 1-palmitoyl-2-oleoyl-sn-glycero-3-phosphatidylcholine
POPE: palmitoyl-2-oleoyl-sn-glycero-3-phosphatidylethanolamine
POPG: 1-palmitoyl-2-oleoyl-sn-glycero-3-phosphatidylglycerol
RMSD: root mean-squared deviation
sTeLIC: ligand-gated ion channel of the endosymbiont of *Tevnia jerichonana*;
TMD: transmembrane domain
ΔV_M_: applied transmembrane potential
XRC: X-ray crystallography

## Author Contributions

M.J.A. and W.W.C designed the electrophysiology studies. M.J.A. expressed and purified WT ELIC and mutants as well as incorporated channels into liposomes. M.J.A. and E.J.W. performed patch clamp experiments and analyzed current traces. M.J.A., J.H., and G.B. designed molecular dynamics simulations. M.J.A. performed molecular dynamics simulations and analyzed trajectories. M.J.A. wrote manuscript. All authors read and edited manuscript.

## Declaration of Interests

The authors of this manuscript declare no competing interests.

## Acknowledgments

The authors also wish to thank the National Institutes of Health (K08-GM152844 to M.J.A. and R35-GM137597 to W.W.C.), the National Science Foundation (NSF DGE 2152059 to G.B.), the Agence Nationale de la Recherche through LABEX DYNAMO (ANR-11-LABX-0011 to J.H.), and the Foundation for Anesthesia Education and Research (MRTG-02-15-2022-Arcario to M.J.A.) for the generous support which enabled these studies. This work used Delta at the National Center for Supercomputing Applications (University of Illinois at Urbana-Champaign) through allocation BIOP240179 from the Advanced Cyberinfrastructure Coordination Ecosystem: Services & Support (ACCESS) program, which is supported by U.S. National Science Foundation grants #2138259, #2138286, #2138307, #2137603, and #2138296.

## Data Availability

All data shown in this paper are available in a public repository (10.5061/dryad.fxpnvx16k) with a CC0 license. MD simulation trajectories are downsampled for portability, but full-length trajectories are available upon request. The in-house analysis scripts will also be housed in the same repository. Plasmid DNA generated in this study is available upon request.

## Notes

### Competing Interest Statement

The authors have declared no competing interest.

